# Arm and head domain in highly conserved lipoprotein modification enzyme Lgt determine functional diversity among bacterial pathogens

**DOI:** 10.1101/2025.03.27.645668

**Authors:** Simon Legood, Ana Oliveira Paiva, Najwa Taib, Tristan Ruffiot, Simonetta Gribaldo, Ivo Gomperts Boneca, Nienke Buddelmeijer

## Abstract

Lipoproteins are important components of the bacterial cell envelope that itself is an excellent target for antibiotics. The post-translational lipoprotein modification pathway is conserved in bacteria in which prolipoprotein phosphatidylglycerol diacylglyceryl transferase (Lgt) catalyses the first and committed step. Due to its essentiality for cell viability in proteobacteria its membrane localisation and relative accessibility, Lgt is proposed as promising target for the development of novel antibiotics. To answer the question of the degree of conservation between Lgt homologues of WHO-listed pathogenic species we performed evolutionary, structural and functional analyses. Our data show that Lgt is present in all bacteria and absent from archaea. Alpha Fold structural models are similar to the X-ray structure of Lgt from *E. coli* with most variability and less conserved residues in the arm- and head domains. Lgt of proteobacteria but not of firmicutes restore growth and viability of a Lgt depletion strain in *E. coli*. Sequence alignments and site-directed mutagenesis demonstrate that unique conserved residues on arm-2 together with histidine 103 determine protein substrate specificity. This large-scale analysis led to the definition of a 13-residue Lgt motif and an alternative catalytic mechanism. Our results highlight similarities in catalytic mechanism and differences in substrate specificity between Lgt homologues from pathogenic species with impact on strategies to develop narrow-spectrum antibiotics targeting Lgt.

**Author summary:** Antimicrobial resistance is a major threat to public health for which novel targets to develop new therapies is urgently needed. The bacterial lipoprotein modification pathway is promising for exploration of new antibiotics since it is unique to bacteria, it is essential for bacterial viability and virulence, and it is accessible to drugs due to the exposed domains of the modification enzymes. In this study we explored large-scale sequence analysis, structural modelling and functional assays of the first enzyme in the pathway. Our findings show that the enzyme is highly conserved across distant phyla, that homologous enzymes have similar structures and contain a signature motif composed of invariant essential residues, but functional conservation divides monoderm and diderm pathogenic bacteria. This correlates with structural variation and differences in substrate specificity, illustrating the potential for the development of narrow spectrum antibiotics targeting the lipoprotein modification pathway.

## Introduction

The bacterial cell envelope fulfils an important function in bacterial physiology by maintaining cell shape and protecting against external threat and environmental stress (1). It also plays an important role in pathogenic species through interactions between its virulence factors and host cells leading to signalling of the innate immune response (2). Lipoproteins are important components of the cell envelope of mono- and diderm bacteria, where they are anchored into phospholipid membranes via their lipid N-terminus (3). Lipoproteins with a classical cysteine-containing recognition sequence (so called lipobox) undergo post-translational modification through a sequential action of three integral membrane enzymes. In the canonical pathway, well characterised in *Escherichia coli*, preprolipoprotein phosphatidylglycerol diacylglyceryl transferase (Lgt) is the first enzyme that adds a diacylglyceryl (DAG) moiety from phosphatidylglycerol onto the cysteine resulting in a thioether linked prolipoprotein (4). Signal peptidase II (Lsp) cleaves the signal peptide (5), followed by apolipoprotein N-acyltransferase (Lnt) that transfers a fatty acid from *sn*-1 of phosphatidylethanolamine resulting in a triacylated mature lipoprotein (6). The lipoprotein modification pathway is largely conserved and Lgt and Lsp are essential for viability in proteobacteria. Although the first two steps are thought to be consistent in all bacteria, variation occurs in the third step of modification. Lnt is essential for viability in γ-proteobacteria but not in other proteobacteria (7–10). This includes ε-proteobacterium *Helicobacter pylori* in which Lnt was shown important for virulence (11). N-acylation varies between species. In *Bacteroides* N-acylation is catalysed by Lnb (12), which is structurally and catalytically different from Lnt of *E. coli* (13). Monoderm bacteria also perform N-acylation of lipoproteins (14) by the combined action of two enzymes LnsA and LnsB (15). These bacteria can also transfer fatty acids from S-diacylglyceryl cysteine to the free α-amine group of the same residue via lipoprotein intermolecular transferase (Lit) (16, 17). The impact of fatty-acid acylation of lipoproteins on signalling of the immune system through interaction with Toll-like receptor 2 heterodimers has been reported (18, 19). In monoderm bacteria, where lipoprotein modification is not essential for viability, the degree of lipidation results in attenuation or enhanced virulence depending on the species (20). Surface exposed lipoproteins in diderm bacteria are potent virulence factors and are developed (8, 21) or explored as vaccine candidates (22).

The lipoprotein modification pathway is considered an excellent target for the development of novel antibacterials (22, 23). Indeed, Lsp inhibitors, globomycin (24) and myxovirescin (25), have been known for many years and the structure of the enzyme-inhibitor complex with globomycin has been solved (26). More recently, two inhibitors of Lgt were identified by selecting cyclic peptides that bind Lgt (27) and in a high-throughput screen with small molecules that inhibit *in vitro* growth of *E. coli* (28). Both cyclic peptide G2428 and inhibitor MAC-0452936 inhibit Lgt activity *in vitro*. The X-ray crystal structure of Lgt from *E. coli* depicts a compact integral membrane enzyme composed of seven transmembrane domains (body), two arms that align on top of the cytoplasmic membrane and a periplasmic domain (head) (29). A central cavity contains essential residues where the diacylglyceryl transfer reaction is predicted to take place. A catalytic mechanism is proposed with a central role for histidine 103 together with residues in the central cavity that interact with the polar headgroup of phosphatidylglycerol (30, 31). Binding of preprolipoprotein is proposed to induce a conformational change in Lgt such that histidine 103 moves close to the phospholipid and the protein substrate in the central cavity where it acts as a catalytic base and activates cysteine in preprolipoprotein for a nucleophilic attack on C3 of phosphatidylglycerol (29, 31).

In the work described here, we asked to what degree Lgt is conserved on an evolutionary, structural and functional level in bacteria, particular pathogenic species that pose a serious threat to human health. Our results show that specific amino acids in a structurally highly conserved protein determine functional variability between Lgt homologues. This suggest that Lgt is suitable for the development of narrow-spectrum antibiotics.

## Results

### Primary sequence and phylogenetic analysis show high conservation of Lgt proteins across distant phyla

We searched Lgt protein homologs in 22 bacterial species listed as priority pathogens (32–35) using the SyntTax database (36). The retrieved sequences were aligned, and a maximum likelihood phylogeny was inferred to obtain insight into the degree of evolutionary conservation. Overall, the multiple sequence alignment (MSA) showed poor conservation in the arms and periplasmic head domains (Fig. 1). It revealed 16 conserved residues including amino acids postulated to be involved in catalysis (29, 31). Thirteen residues are essential for viability (29, 30, 37, 38) among which histidine 103 found in proteobacteria but not in firmicutes and actinobacteria where Y and W substitutes H103, respectively. Lgt of *M. tuberculosis* has a large C-terminal extension that has no homology with known protein sequences. Based on computational modelling H103 is proposed to activate the thiol group of the cysteine residue of preprolipoprotein leading to transfer of the diacylglyceryl (DAG) moiety from phosphatidylglycerol (31).

**Fig. 1.**
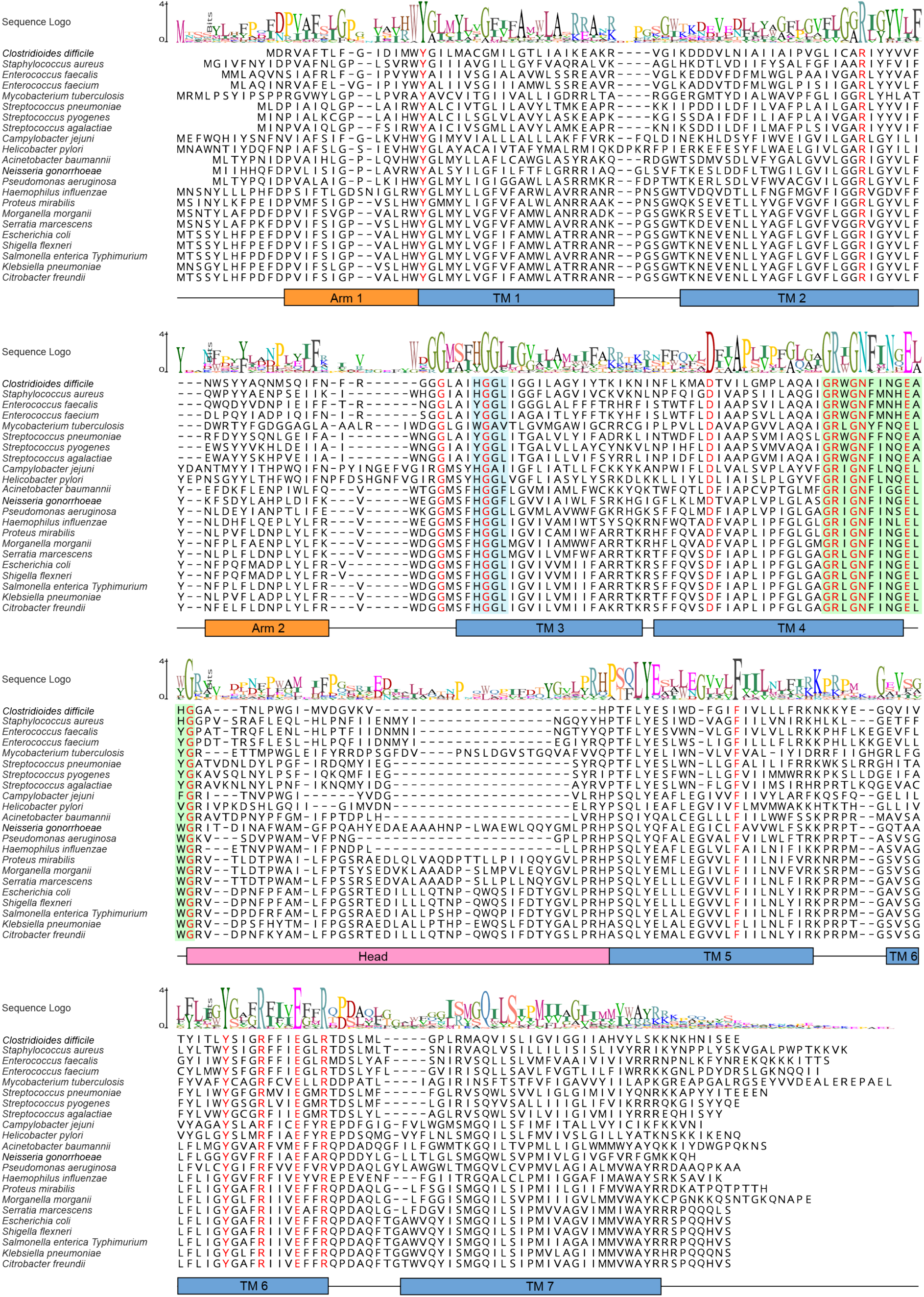
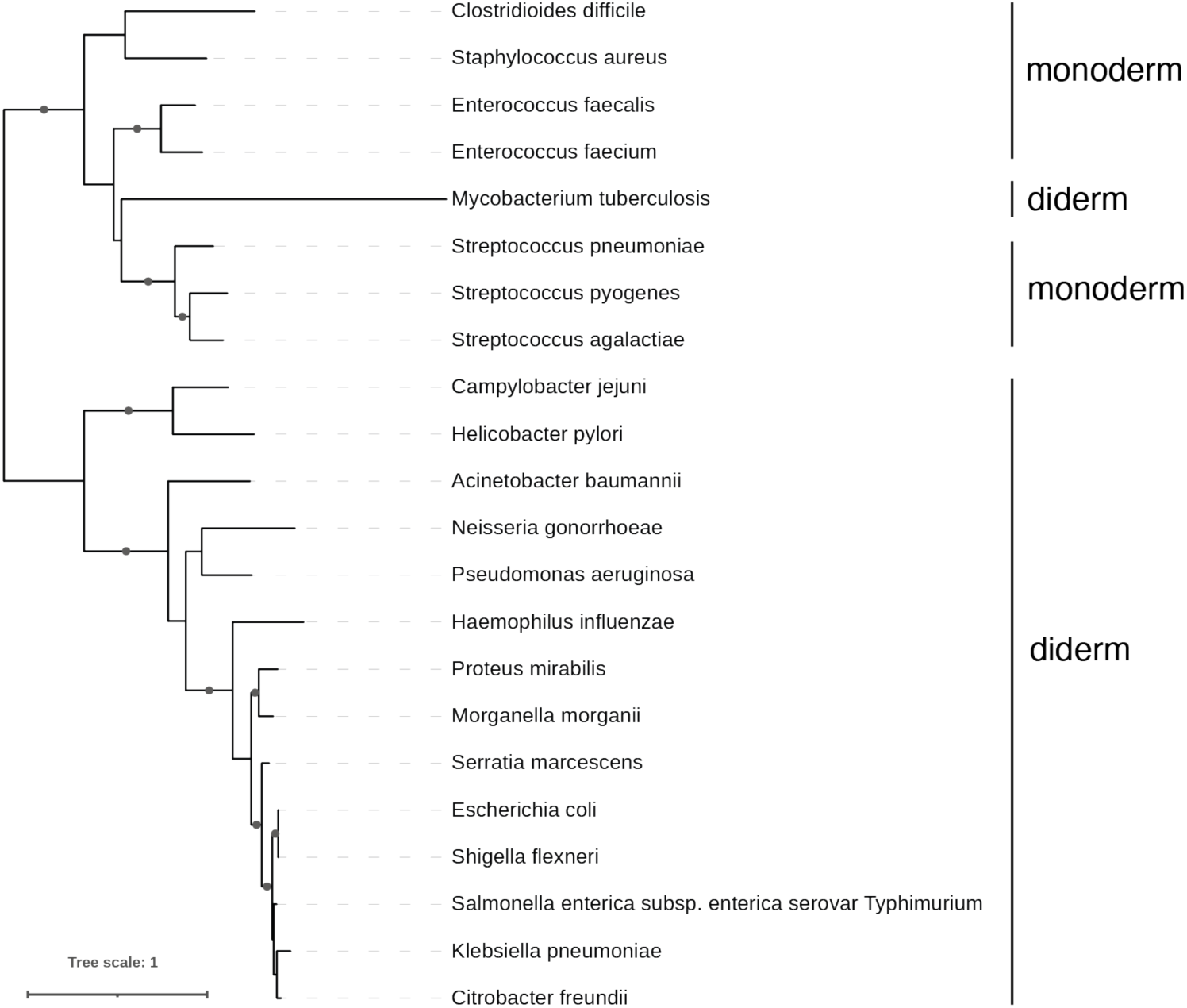
Sequence alignment and phylogenetic tree of Lgt proteins. (A) Twenty-two Lgt proteins were selected from bacterial pathogens. Degree of amino acid conservation is indicated in sequence logo with 16 fully conserved residues shown in red, including potential catalytic residues R143, N146, Y235 and R239 (*E. coli* numbering). Conserved motif HGGL (blue) containing active site residue H103 (W/Y) and the Lgt signature motif (green) are highlighted. Transmembrane domains TM1-TM7 (blue), periplasmic head domain (corresponding to Β4-Β5-α3-α4-Β6) (pink) and arm-1 (Β2-Β3) and arm-2 (α1-ρι2) (orange) are shown according to the X-ray structure of *E. coli* Lgt (29). The Lgt sequence of *M. tuberculosis* is truncated at the C-terminus at position 303. (B) The maximun-likelihood tree was inferred from an alignment of 22 sequences and 222 amino acid positions with the model LG+F+G4. The scale bar represents the average number of substitutions per site. Black dots indicate bootstrap values >90%.

The Lgt phylogenetic tree has two distinct clades, the first containing Lgt sequences from the phylum firmicutes and the second containing proteobacteria Lgt sequences (Fig. 1). Lgt of *Mycobacterium tuberculosis*, an actinobacterium, branches within lactobacillalles. The separation between proteobacteria and firmicutes, was previously observed but not further explored, where Lgt of *M. smegmatis* branches between proteobacteria and firmicutes (39). These results suggest a vertical inheritance of Lgt in bacteria.

Next we analysed the genomic context of *lgt* in the 22 species (36). Synteny is observed for all species except for Lgt of *M. tuberculosis* and the genomic context is only conserved in enterobacteriales (Suppl. Fig. 1). So, while the Lgt sequences are conserved, the genomic organisation surrounding *lgt* is not.

### The Lgt structure is highly conserved with main variations in the periplasmic (head) domain

Two X-ray crystal structures of Lgt from *E. coli* were solved, form-1 (PDB 5azb) with palmitate and detergent β-octylglucoside, and form-2 (PDB 5azc) with phosphatidylglycerol and DAG referred to as the active form of the enzyme (29). The essential and conserved residues reside on top of TM-3 and 4 and in TM-5 and 6, the exposed periplasmic (head) domain is least conserved (Fig. 1).

Structural conservation of Lgt homologs was assessed using structure prediction program AlphaFold2 (AF2) (40). Confidence metrics for Lgt of *E. coli* state high confidence by pIDDT and low prediction error as defined by PAE, however, for the region at the periplasmic half of TM-7 and the loop between TM-6 and 7 (L6-7) low pIDDT and high PAE scores were observed suggesting flexibility in this region (Suppl. Fig. 2). Comparison of the AF2 structure and X-ray structures clearly shows the difference in this region, i.e. the X-ray structure has an extended α-helix that arcs below the membrane plain whereas the predicted AF2 model has a shortened α-helix and unstructured loop extending upwards through the membrane (Fig. 2).

**Fig. 2.**
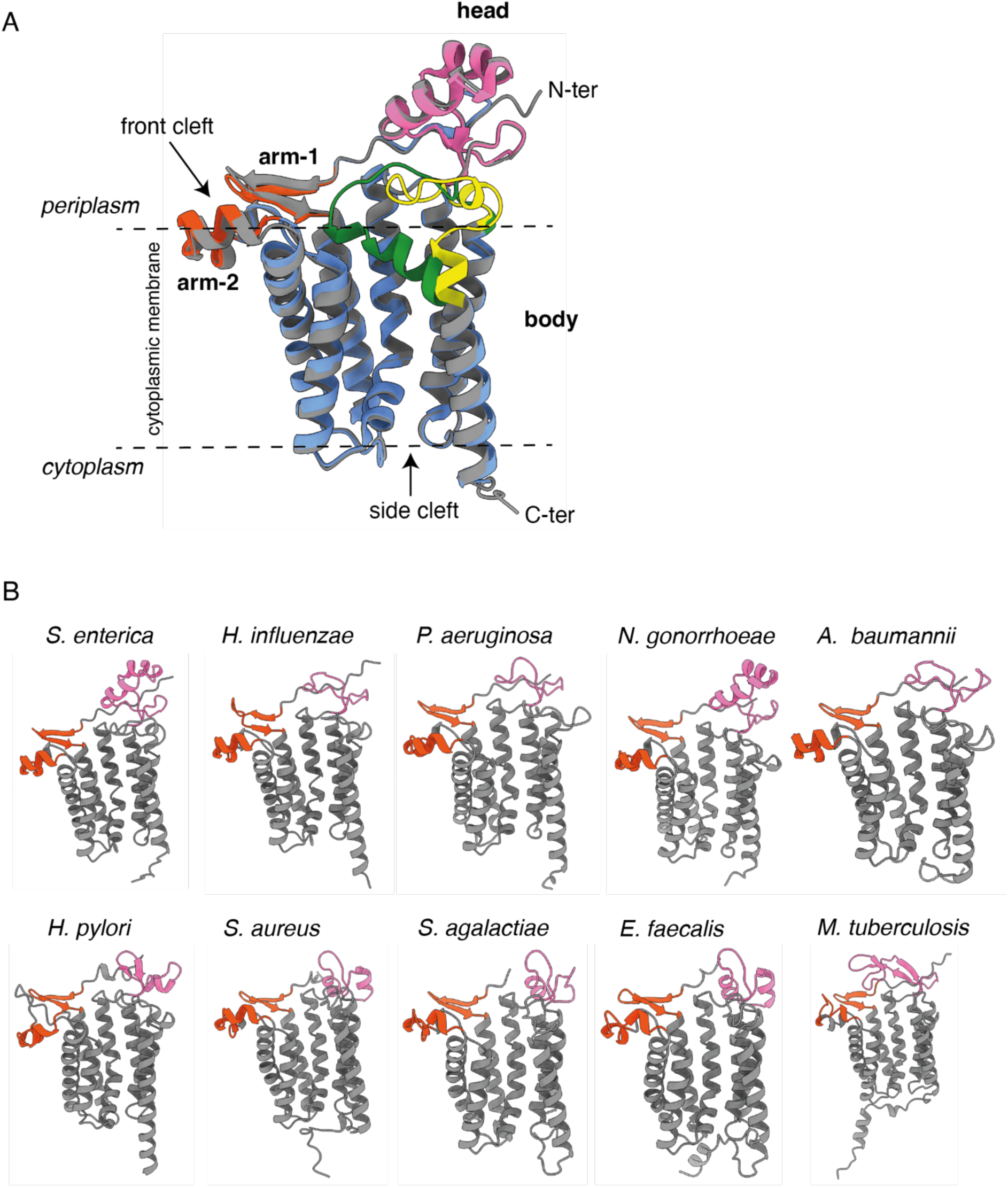
Lgt X-ray structure and Alpha Fold 2 models from selected pathogenic species. (A) Comparison of X-ray structure PDB 5azc (blue) and AF2 structure model of *E. coli* Lgt (grey). Difference in conformation of TM7 and loop connecting TM6 and 7 between X-ray structure and AF2 model are indicated in green and yellow, respectively. Entry of substrate(s) is predicted to occur at the front cleft formed by arm-1 and arm-2 (shown in orange), exit of diacylglyceryl prolipoprotein is predicted via the side cleft (31). The head domain (pink) is exposed to the periplasm. (B) AF2 structural models of 10 Lgt proteins of pathogenic species as shown in Fig. 1. Arm domains are shown in orange and head domain in pink. The C-terminus of Lgt from *M. tuberculosis* is truncated at position 303.

From the 22 Lgt proteins, 10 were selected based on pathogen priority profile and degree of antibiotic resistance and modelled using AF2 (Fig. 2). Due to an extension at the C-terminus of Lgt of *M. tuberculosis*, low confidence and high prediction error were calculated. Main differences among models were observed in L6-7 and in the loop between arm-2 and TM-3 (Fig. 2). The most striking difference in structure, consistent with poor primary sequence conservation, is found in the periplasmic (head) domain. Enterobacteriales and β-proteobacteria have a similarly large head domain whereas the more distant related proteobacteria and firmicutes have smaller less prominent head domain that are predicted to be located closer to the membrane interface (Fig. 2). These results suggest that although Lgt proteins are highly conserved and catalyse the same diacylglyceryl transfer reaction, they may differ in enzymatic activity and/or substrate specificity.

### Lgt of proteobacteria but not firmicutes are functional in *E. coli*

We sought to determine the degree of functional conservation of Lgt from different species by employing complementation studies using the Lgt depletion strain of *E. coli* Δ*lgt* (27, 41). The endogenous *lgt* gene is replaced by a kanamycin resistance cassette and *lgt* is expressed under control of an arabinose inducible promoter (P*_ara_* inserted at the *λ*-attachment site) (27). We previously described a low copy expression vector containing *lgt* genes under control of a P*_lac_* promoter for use in complementation studies (38, 41). Genes encoding Lgt of species representing different branches of the phylogenetic tree were cloned in this vector to study their function in *E. coli* (Suppl. Table 1).

We performed growth kinetics and cell viability analyses to determine the capacity of Lgt homologues to restore growth in liquid medium and to form colonies, respectively. When chromosomal *lgt* expression was repressed by growth in D-glucose and homologous *lgt* expressed by induction with IPTG, viability was not restored in presence of empty vector (v) (Fig. 3). Colonies formed with Lgt of *E. coli*, *Haemophilus influenzae*, *Helicobacter pylori* and in lower numbers with Lgt of *Pseudomonas aeruginosa*, but not with Lgt from *Neisseria gonorrhoeae*, *Salmonella enterica* serovar Thyphimurium and *Acinetobacter baumannii*. Complementation by Lgt of *H. pylori* results in a small colony phenotype on plates (Fig. 3).

**Fig. 3.**
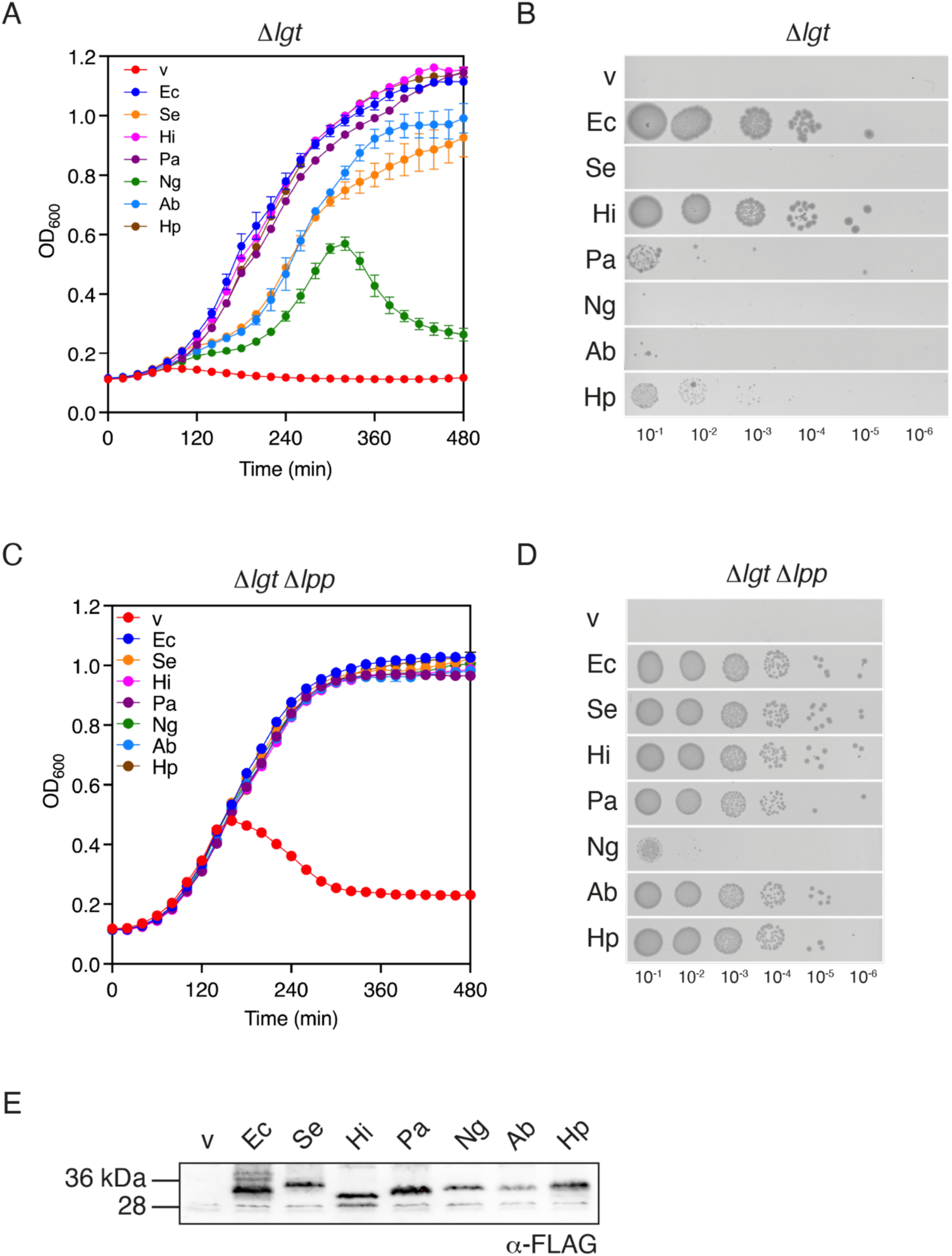
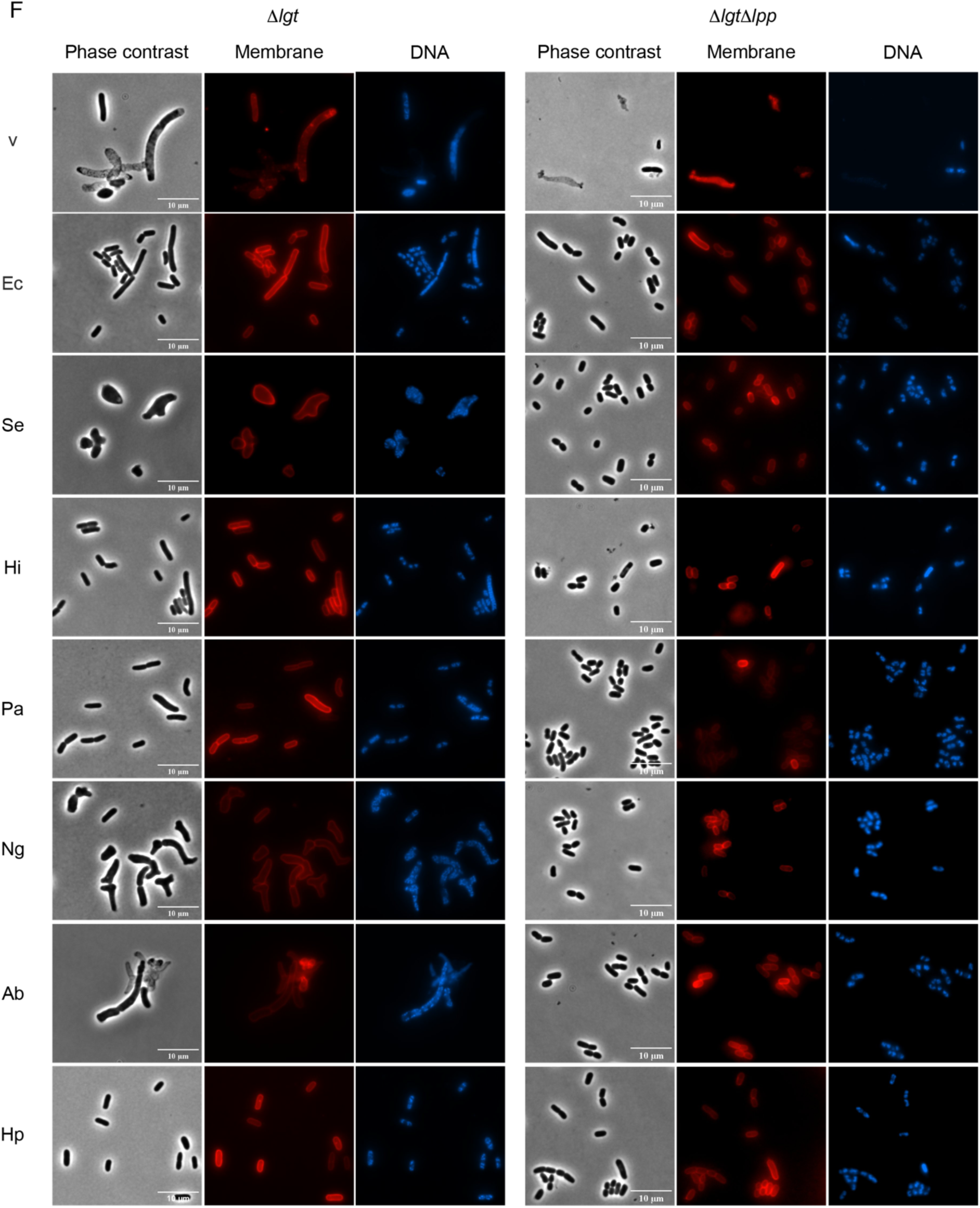
Lgt from proteobacteria complement an *E. coli* Lgt depletion strain in absence of major lipoprotein Lpp. (A) Growth of Δ*lgt* (SLEC67) carrying low copy plasmid pAM238 encoding Lgt homologs in 0.2% D-glucose and 5 mM IPTG. After initial depletion of Lgt through growth in absence of sugar, cultures were grown in 96-well plates in a TECAN spectrophotometer and OD_600_ was recorded at regular intervals. Assays were completed at least in duplicates; error bars indicate standard deviation from the mean. v: empty vector; Ec: *E. coli*; Se: *S. enterica* serovar Thyphimurium; Hi: *H. influenza*; Pa: *P. aeruginosa*; Ng: *N. gonorrhoeae*; Ab: *A. baumannii*; Hp: *H. pylori*. (B) Spot dilution assay of strains grown in the presence of 0.2% D-glucose and 5 mM IPTG corresponding to T_0_ of the growth curve. Assays were completed in duplicate. (C) Growth of Δ*lgt*Δ*lpp* (SLEC68) carrying low copy plasmid pAM238 encoding Lgt homologs under the same conditions as in panel A. (D) Spot dilution assay of SLEC68 strains grown in the presence of 0.2% D-glucose and 5 mM IPTG corresponding to T_0_ of the growth curve as shown in C. (E) Detection of Lgt-FLAG_3_ upon expression of *lgt-flag_3_* from low copy plasmid pAM238 in *E. coli* MG1655^Q^ by addition of 5 mM IPTG. Total cell lysates corresponding to 0.1 OD_600_ units were loaded to each lane. (F) Microscopy images of fixed cells corresponding to T_270_ of the growth curves. Membranes were stained with lipophilic dye FM4-64X (red) and nucleoids by Hoechst 2333 (blue). Scale bar 10 μm.

Growth is delayed with Lgt from *A. baumannii* and *Salmonella enterica* serovar Thyphimurium and reaches only exponential phase with Lgt of *N. gonorrhoeae*. Cell morphology is strongly affected in all three strains; larger and branched cells can be observed, while complementing Lgt homologs give rise to a normal cell morphology (Fig. 3). Strikingly, while Lgt from *S. enterica* is a close homolog of *E coli* it does not restore morphology and viability of Δ*lgt*. All Lgt homologs are produced in wild type *E. coli* (Fig. 3) and while protein quantity is variable, similar amounts are observed for Lgt of *S. enterica* and *H. influenzae* for example.

We previously showed that lipoprotein Lpp (Braun’s lipoprotein) is restricted to a subclade of γ-proteobacteria (42). Lpp is the most abundant protein in *E. coli*, which when mis-localised to the cytoplasmic membrane and covalently cross-linked to the peptidoglycan leads to cell lysis (43). We recently demonstrated that Lgt is essential for growth and viability of *E. coli* in the absence of Lpp highlighting the importance of other lipoproteins for physiology (41). Due to the high abundance of Lpp, we reasoned that subtle differences in function between Lgt homologs could be more clearly observed in a Lgt depletion strain lacking Lpp (Δ*lgt*Δ*lpp*) (41). Upon induction of gene expression with IPTG, all Lgt proteins from proteobacteria led to colony formation, except with Lgt of *N. gonorrhoeae* (Fig. 3). Growth is fully restored in liquid media in the presence of IPTG, including *N. gonorrhoeae* (Fig. 3). In the absence of Lpp, cell morphology is restored and was similar to cells complemented with Lgt of *E. coli* (Fig. 3). These data suggest that some Lgt homologues from proteobacteria are less efficient in modifying *E. coli* lipoproteins and are functional when substrate load is reduced through elimination of Lpp.

Lgt from *Enterococcus faecalis*, but not from *Staphylococcus aureus* or *Streptococcus agalactiae*, was produced in *E. coli* (Fig. 4). Codon-optimization of *lgt* from *S. aureus* and *S. agalactiae* for expression in *E. coli* slightly increased protein production of *S. agalactiae* Lgt but not of *S. aureus* Lgt (Fig. 4). Lgt from *E. faecalis* and *S. agalactiae* did not result in colony formation or growth in liquid media due to cell lysis of Δ*lgt*Δ*lpp* (Fig. 4), indicating that these proteins are not functional in *E. coli*.

**Fig. 4.**
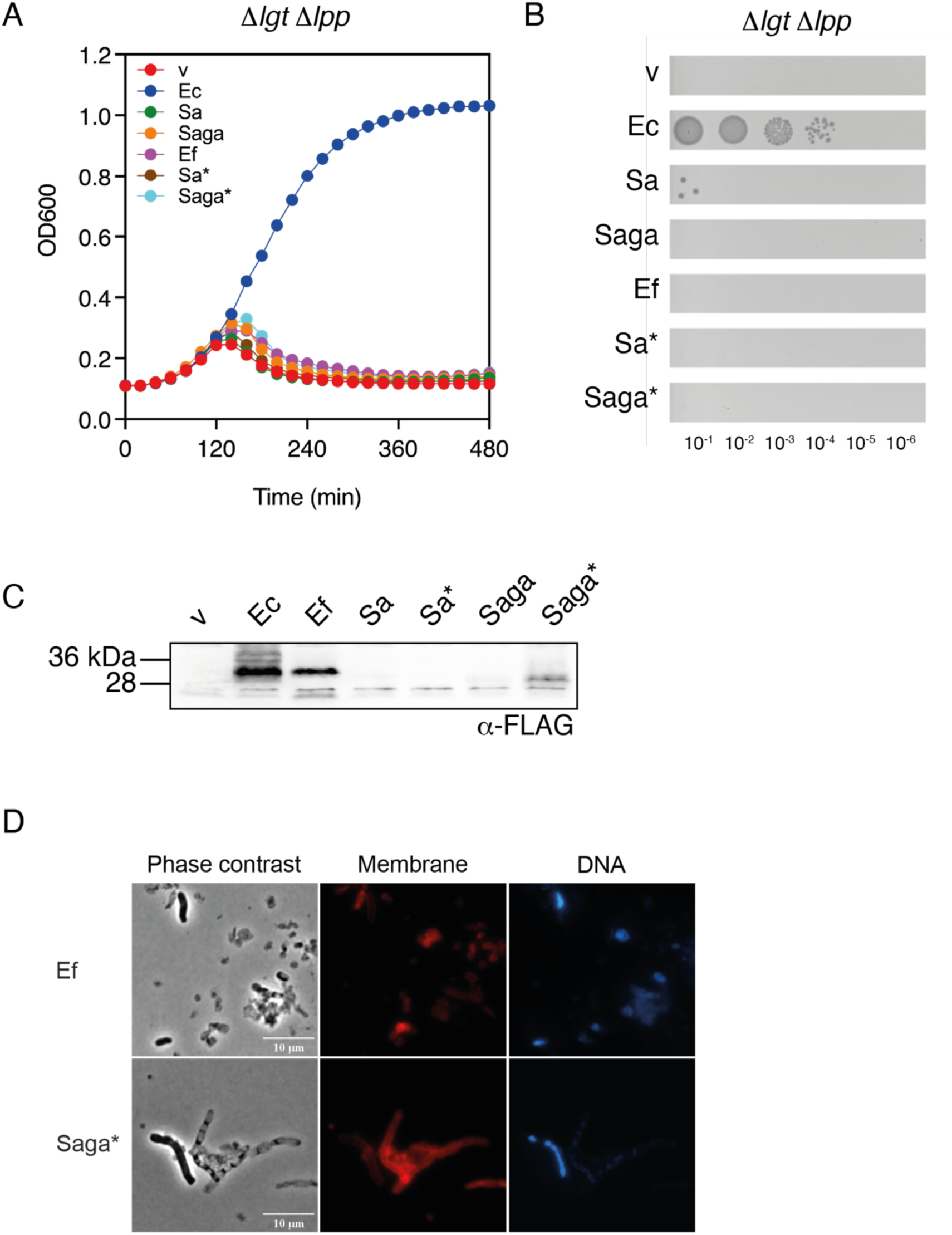
Lgt homologues from firmicutes are not functional in *E. coli*. (A) Growth of Δ*lgt*Δ*lpp* (SLEC68) containing plasmids with *lgt* genes from firmicutes in the presence of 0.2% D-glucose and 5 mM IPTG. Asterisk represents codon-optimized *lgt* genes from *S. aureus* (Sa) and *S. agalactiae* (Saga) for expression in *E. coli*. Ef: *E. faecalis*. Assays were completed in at least duplicates; error bars indicate standard deviation from the mean. (B) Spot dilution assay of strains grown in the presence of 0.2% D-glucose and 5 mM IPTG corresponding to T_0_ of the growth curve shown in panel A. (C) Detection of Lgt-FLAG_3_ upon expression of *lgt-flag_3_* from low copy plasmid pAM238 in *E. coli* MG1655^Q^ by addition of 5 mM IPTG. Total cell lysates corresponding to 0.1 OD_600_ units were loaded to each lane. (D) Microscopy images of fixed cells corresponding to T_270_ of the growth curve. Membranes were stained with lipophilic dye FM4-64X (red) and nucleoids with Hoechst 2333 (blue). Scale bar 10 μm.

Together the results show that the capacity of Lgt proteins to function in *E. coli* reflects the two-way division observed in phylogeny and suggests the presence of domains and/or specific amino acids reflecting differences in enzyme function and/or substrate specificity.

### The periplasmic (head) domain is important for Lgt function

The main differences in sequence and predicted structure among Lgt proteins are the arms and head domain; overall, the head domain of Lgt of firmicutes is smaller than those of proteobacteria (Fig. 2). We constructed so-called head domain swap proteins between *E. coli* Lgt and corresponding domains of Lgt from *H. pylori*, *S. aureus* and *M. tuberculosis* to determine its role in Lgt function. Lgt from *H. pylori* (functional in *E. coli*) and *S. aureus* (not produced in *E. coli*) have small head domains, 17 and 23 residues, respectively. The head domain of Lgt of *M. tuberculosis* (33 amino acids) is similar in size and was selected based on the divergence between firmicutes and proteobacteria (Fig. 1). All three head swap proteins are produced in *E. coli* (Fig. 5). Expression of Lgt-Head^Hp^ in the Lgt depletion strains resulted in colonies, albeit small in Δ*lgt*, and growth in liquid medium as observed with *E. coli* Lgt even in the presence of Lpp. Its morphology is similar to the complemented strain with Lgt of *E. coli* (Fig. 5 and Fig. 3).

**Fig. 5.**
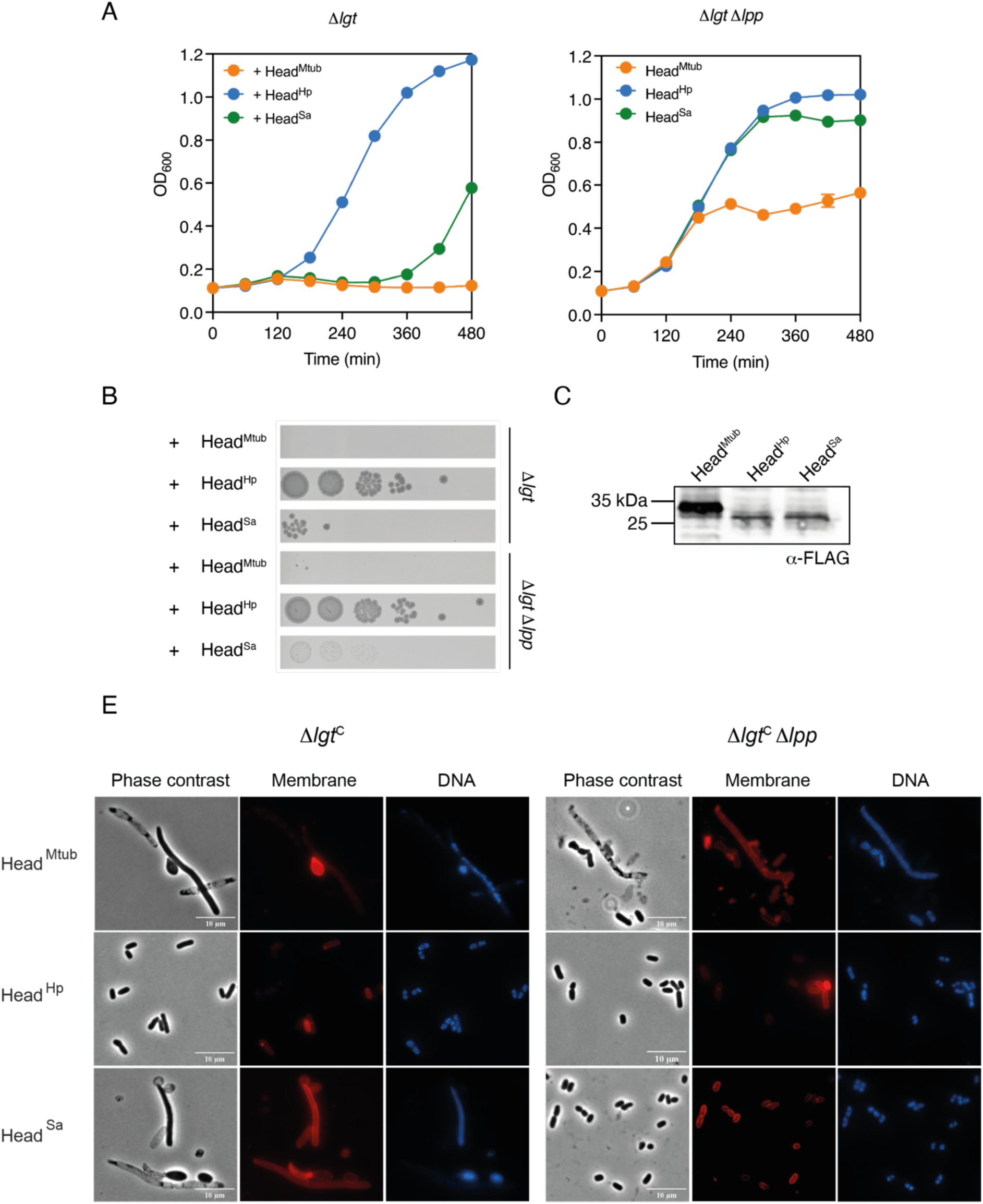
The periplasmic head domain is important for Lgt function. (A) Growth of Δ*lgt* (SLEC67) and Δ*lgt*Δ*lpp* (SLEC68) containing *E. coli lgt* with substituted head domains from *M. tuberculosis* (Head^Mtub^), *H. pylori* (Head^Hp^) and *S. aureus* (Head^Sa^), encoded on low copy plasmid pAM238 in 0.2% D-glucose and 5 mM IPTG. Assays were completed in at least duplicates; error bars indicate standard deviation from the mean. (B) Spot dilution assay grown on plates with 0.2% D-glucose and 5 mM IPTG corresponding to T_0_. (C) Detection of Lgt-FLAG_3_ on Western blot from total cell lysates of MG1655^Q^. Equal cell mass (0.1 OD_600_ units) was loaded per lane. (D) Microscopy images of fixed cells corresponding to T_270_ of the growth curve. Membranes were stained with lipophilic dye FM4-64X (red) and nucleoids with Hoechst 2333 (blue). Scale bar 10 μm.

Colony formation was not observed with Lgt-Head^Mt^ in either Δ*lgt* or Δ*lgt*Δ*lpp* but growth increased to mid-exponential phase in the latter. Filamentous cells and lysis are observed in Δ*lgt* and a mixed population is observed in Δ*lgt*Δ*lpp*. Lgt-Head^Sa^ does not completely restore viability in Δ*lgt* and small colonies are formed on glucose IPTG plates in Δ*lgt*Δ*lpp*. Growth is only observed in Δ*lgt*Δ*lpp*. Cell filamentation and lysis is observed in Δ*lgt* and small rod-shaped cells are observed in the Δ*lgt*Δ*lpp* background. These results suggest that the periplasmic head domain is important for Lgt function.

### Essential residues and their role in Lgt function

Several essential residues have been identified in Lgt of *E. coli*, 16 of which are completely conserved in the 22 Lgt proteins from pathogenic species (Fig. 1). Either a temperature sensitive mutant *lgt*(ts) (30, 37) or a Lgt depletion strain where the arabinose controlled *lgt* is located on a plasmid (29, 38) were used previously in complementation experiments. Here, we took advantage of strain Δ*lgt*Δ*lpp* and our previously reported Lgt variants to obtain a more detailed insight into the role of essential residues in Lgt function. All Lgt proteins are produced in *E. coli*, but quantity varies as previously reported (Fig. 6) (38). Variants with D129A and E243A grow as non-mutated Lgt and are viable. Lgt with G98 located between arm-2 and TM-3, G104 in TM-3, and E151 located on the loop between TM-4 and the head domain, substituted by alanine are delayed in growth. These proteins are produced in slightly higher quantity compared to non-mutated Lgt that might affect efficacy of the reaction. Lgt with Y26A (TM-1), R143A (TM-4), N146A (TM-4), G154A (loop between TM-4 and head domain) and R239A (TM-6) mutations did not restore growth (Fig. 6). Growth of Δ*lgt*Δ*lpp* containing Lgt-H103Q grows to mid exponential phase but optical density does not decrease rapidly. The non-functional Lgt variants and H103Q are not viable and cause cell lysis (Fig. 6). These results confirm the essentiality of Y26, H103, R143, N146, G154 and R239 in Lgt of *E. coli* (29, 38). Four glycine residues are identified among the important amino acids. Their position close to potential catalytic residues H103 (G104), R143 (G142) and N146 (G145) suggest a role in conformational changes upon substrate binding (31, 44). G154 is located between body and head domains (Fig. 7).

**Fig. 6.**
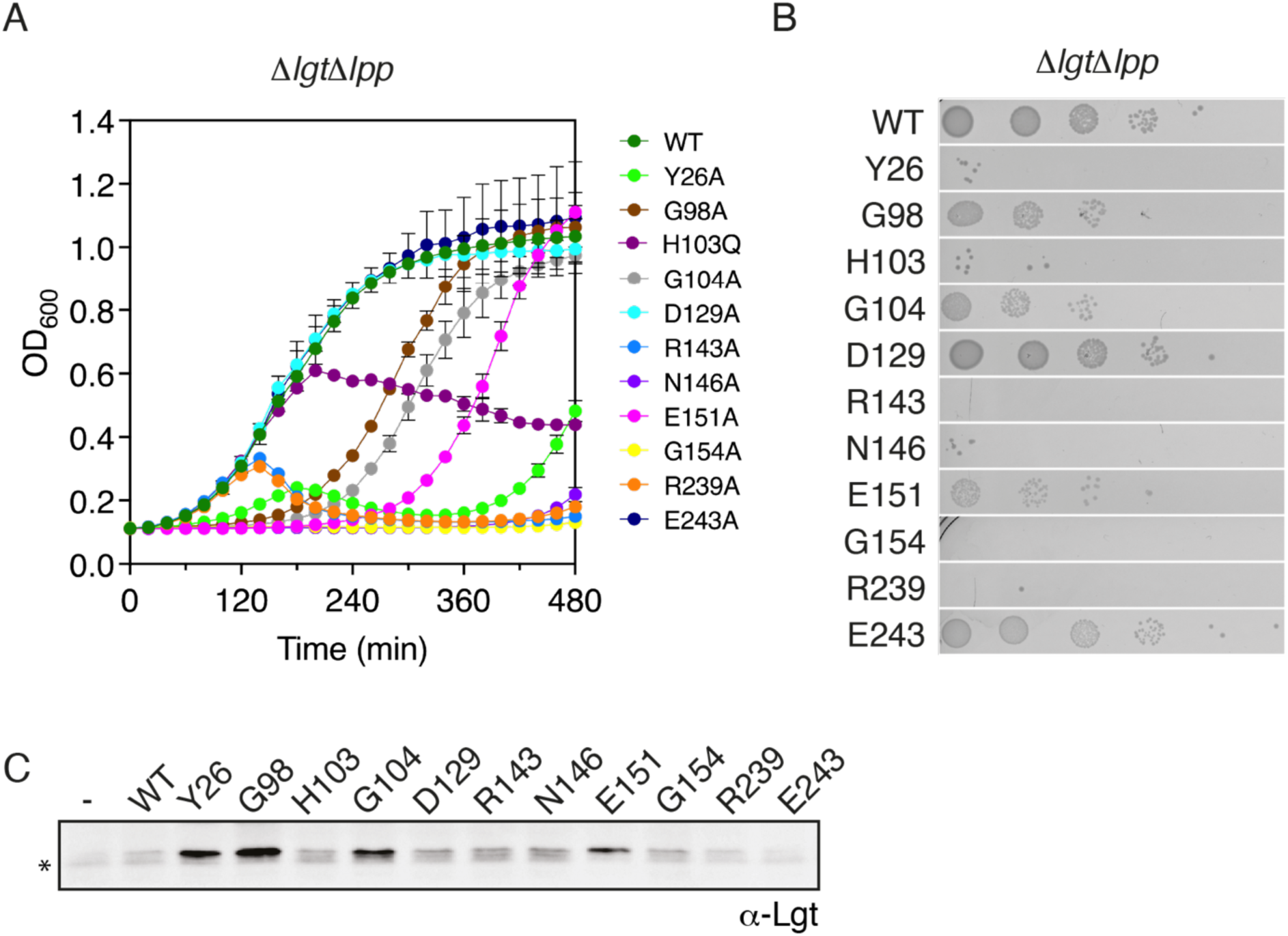
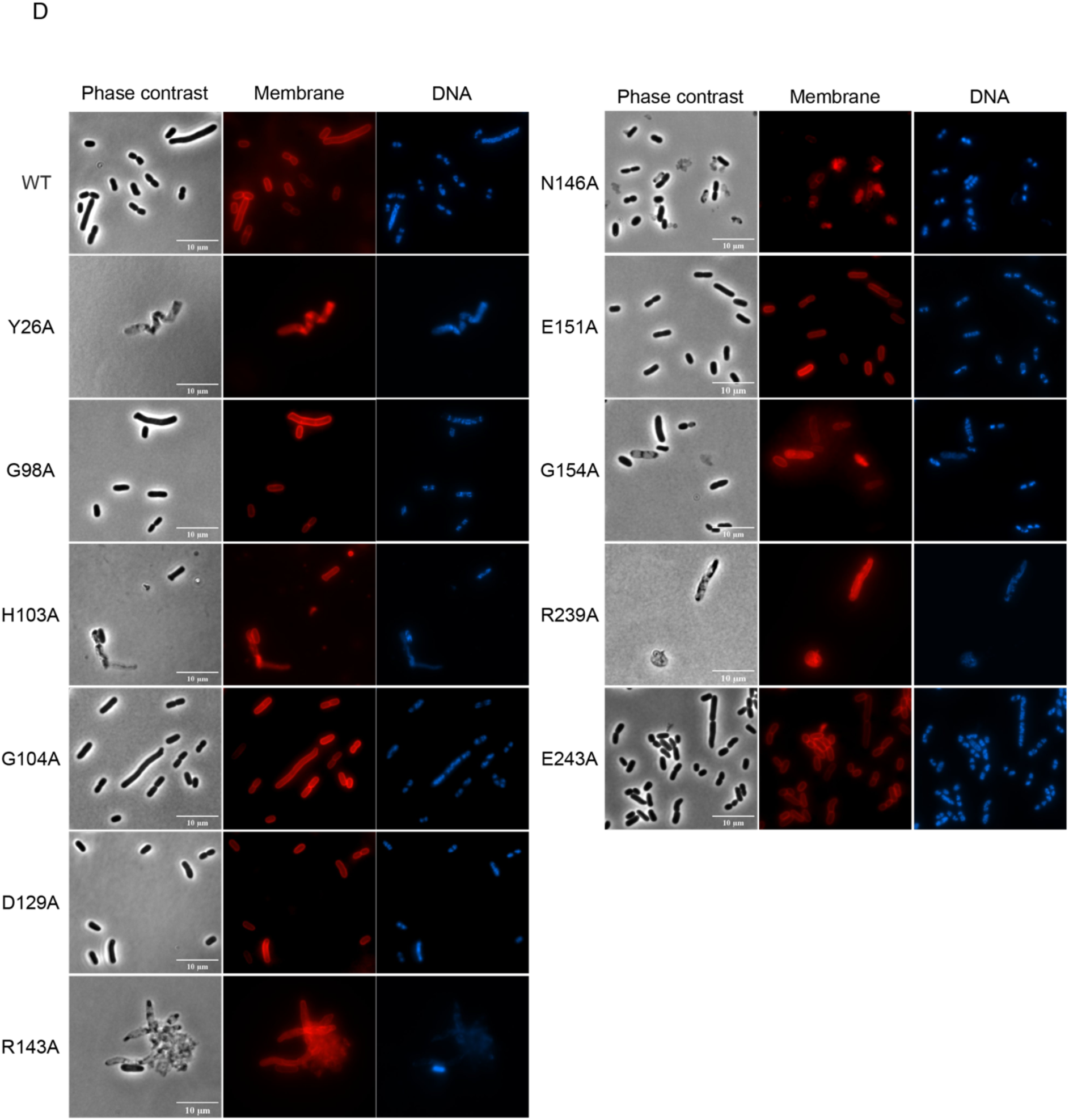
Functional characterisation of essential residues in Lgt of *E. coli*. (A) Growth of Δ*lgt*Δ*lpp* (SLEC68) with pAM238 plasmids expressing alanine substitution mutants and H103Q mutant of Lgt of *E. coli* (38) in 0.2% D-glucose and 5 mM IPTG. (B) Spot dilution assay grown on plates with 0.2% D-glucose and 5 mM IPTG corresponding to T_0_. Assays were performed in duplicate. (C) Western blot detection of Lgt-MYC_2_ proteins in total cell lysates of MG1655^Q^ with anti Lgt antibodies. Asterisk indicates chromosomal Lgt. (D) Microscopy images of fixed cells corresponding to T_270_ of the growth curve. Membranes were stained with lipophilic dye FM4-64X (red) and nucleoids with Hoechst 2333 (blue). Scale bar 10 μm.

**Fig. 7.**
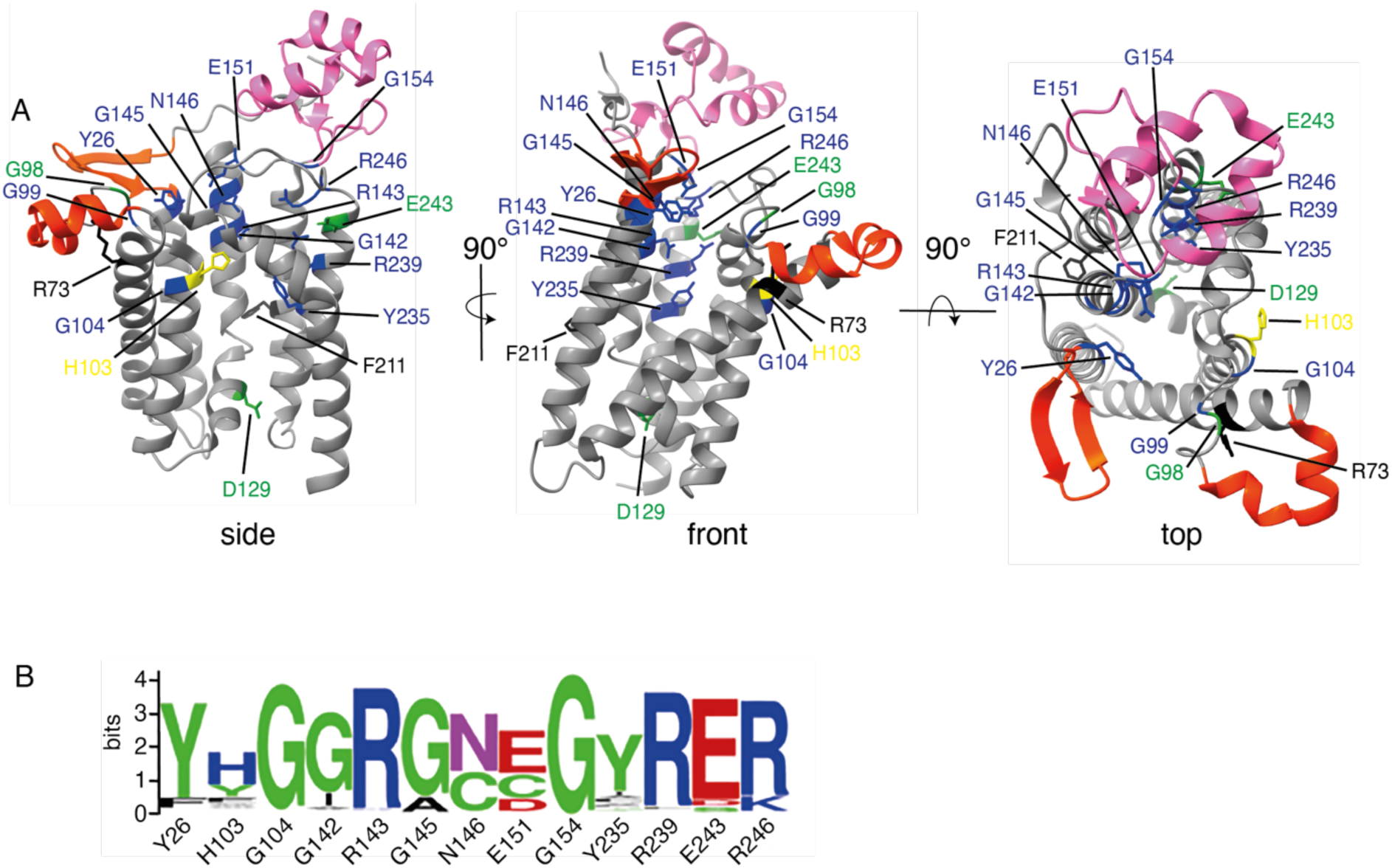
Structural conservation of proposed catalytic- and essential residues in Lgt of proteobacteria. (A) Conserved residues are depicted on X-ray crystal structure 5azc of *E. coli* Lgt. H103 (yellow) is essential but not completely conserved among bacteria and can be replaced by Y or W (Fig. 1). Essential residues are shown in blue, non-essential residues (D129 and E243) in green. Conserved residues R73 and F211 have not been described (black). (B) Essential amino acids composing the 13 residue Lgt motif as WebLogo.

We next identified residues in Lgt that are unique for proteobacteria and absent in Lgt from firmicutes to obtain clues about amino acids implicated in Lgt function other than catalysis that may determine substrate specificity (Suppl. Fig. 3). We performed MSA of the 14 Lgt-s from proteobacteria and 7 Lgt-s from firmicutes separately and identified the conserved residues in each group. Out of 22 residues that are unique in proteobacteria 5 amino acids are essential. Y30, Y80, M100, and S101 are located at the front cleft close to arm-2, and G263 is located in the bend of TM-7 (Suppl. Fig. 4). These results suggest that entry of substrates through the front cleft is guided by arm-2 via interaction with specific residues that determine substrate specificity and bending of TM-7 by G263 provides dynamics for catalysis.

### Lgt of Bacteriodes thetaiotaomicron is not functional in *E. coli*

One of the major challenges in antimicrobial development is to protect the microbiota when treating bacterial infections caused by pathogenic species. *Bacteriodes thetaiotaomicron* (*B. theta*) is an obligate anaerobic diderm bacterium and prominent member of the human gut microbiota (45). Sequence alignment of Lgt from *B. theta* with Lgt of *E. coli* shows low conservation although an AF2 structural model resembles Lgt of *E. coli* as seen for other Lgt proteins (Suppl. Fig. 4). Three amino acids changes are observed among the 16 conserved residues in *B. theta* Lgt, i.e. G142I, Y235I and R246K. Lgt of *B. theta* does not restore growth of Δ*lgt*Δ*lpp* in liquid medium due to cell lysis and does not form colonies on glucose/IPTG plates, indicating that it is not functional in *E. coli* (Fig. 8).

**Fig. 8.**
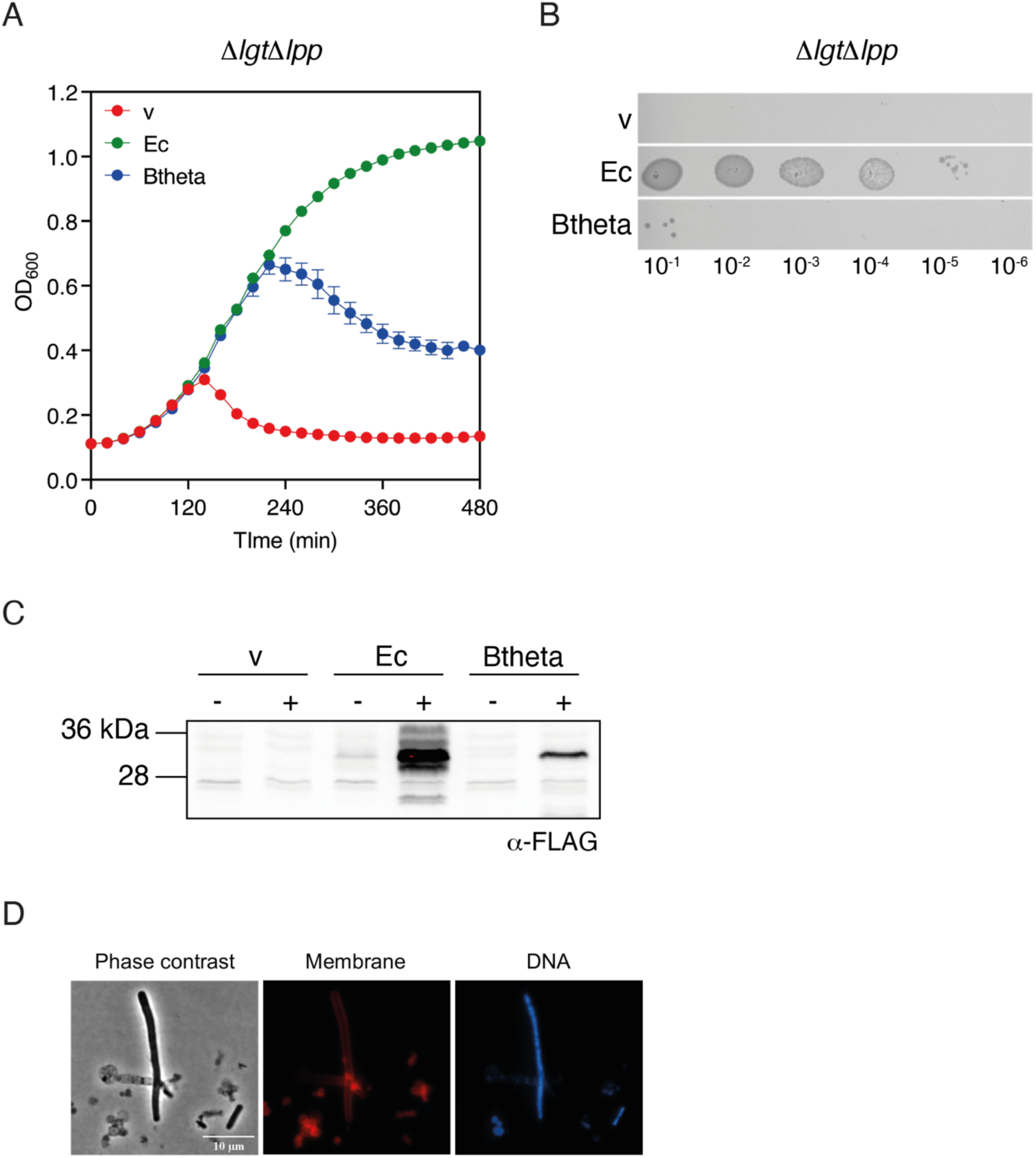
Lgt from *B. thetaiotaomicron* is not functional in *E. coli*. (A) Growth of Δ*lgt*Δ*lpp* (SLEC68) expressing *lgt* from *B. thetaiotaomicron* from P*_lac_* on pAM238 in 0.2% D-glucose and 5 mM IPTG. (B) Spot dilution assay on plates with 0.2% D-glucose and 5 mM IPTG corresponding to cultures of T_0_. (C) Western blot of Lgt-FLAG_3_ in MG1655^Q^ lysates grown in the absence (-) and presence (+) of 5 mM IPTG. (D) Microscopy images of fixed cells corresponding to T_270_ of the growth curve. Membranes were stained with lipophilic dye FM4-64X (red) and nucleoids with Hoechst 2333 (blue). Scale bar 10 μm.

### Lgt is evolutionary conserved in all bacteria and absent from Archaea

We next applied a phylogenomic approach to explore the conservation of Lgt at a larger evolutionary scale and the molecular mechanism of the reaction. We first searched the refseq_genomes database for prolipoprotein phosphatidylglycerol diacylglyceryl transferase encoding genes using the Pfam domain PF01790. At the time of analysis (02/2021) 14664 hits were identified in 12956 out of 13512 different genomes (Suppl. Table 2). Lgt is found in bacteria, including those that lack a peptidoglycan cell wall, and is absent from archaea, although the presence of lipoproteins has been reported (46, 47). Genomes with more than one Lgt tend to be larger than average; there are approximately 1300 genomes with two genes and less than 200 with up to four copies. Several species possess two *lgt* genes that confirm literature reports, as for example *Streptomyces coelicolor* (48) and *M. smegmatis* (49), and in members of firmicutes (for example the *B. cereus* group) of which one *lgt* gene is frequently located on a plasmid (50). Plasmids with a copy of *lgt* also encode lipoproteins implicated in DNA conjugation via a type IV secretion machinery. Some species have up to four *lgt* genes (*Deinococcus,* halophilic *Marinilactibacillus*), bacteria with small genomes lack a *lgt* gene, for example endomutualists (Buchnera) or endopathogens (Orientia).

To obtain more detailed information on conserved domains and residues we analysed 179 Lgt sequences from 160 genomes covering all phyla (42) of which 45% were annotated in the refseq_genome database. Lgt from *E. coli* K12, *H. pylori* 26695 and *M. tuberculosis* H37Rv were included as reference sequences. Phylogenetic data show that essential residues in Lgt are highly conserved (Suppl. Fig. 5).

Based on complementation results reported here and, in the literature, we propose a Lgt motif of 13 conserved residues, including proposed catalytic residues: Y_26_ H_103_ G_104_ G_142_ R_143_ G_145_ N_146_ E_151_ G_154_ Y_235_ R_239_ E_243_ R_246_ (Fig. 7). In most species this motif is conserved with some variations at H103 that is replaced by Y, W, or F, even in closely related bacteria, H103 is thus not absolutely conserved. In more distantly related Lgt-s N146 and E151 are replaced by CD or CC (Suppl. Fig. 5). In *M. smegmatis*, one Lgt protein (MSMEG_3222) is the major diacylglyceryl transferase and contains N146, while the second homologue (MSMEG_5408) possesses Y26H and N146C with presumed low Lgt activity (49). Finally, one subgroup contains amino acids that greatly vary from the conserved Lgt motif and even lack the fully conserved R143 and R239 residues (Suppl. Fig. 5). These include *Deinococci*, *Acidimicrobiales*, *Anaerolineales* and *Acidobacteriales* and *Myxococcales*. The absence of these residues, which are proposed to bind the glycerol headgroup of the phospholipid and perform a gating function (R239) (31), suggests that these Lgt enzymes perform a diacylglyceryl transferase reaction through a different mechanism.

In conclusion, specific residues located at the front cleft allow entry of selective substrates. Catalysis occurs in the central cavity by a conserved mechanism that involves binding of phospholipid and activation of the thiol group of cysteine in the lipobox of the preprolipoprotein. The precise catalytic mechanism needs to be further determined but likely involves residues other than H103.

## Discussion

The lipoprotein modification pathway has gained potential as a target for antibacterial development due to the essentiality of the enzymes in proteobacteria and their membrane localisation providing easy access to small inhibitory molecules (22, 23). Lgt catalyses the first committed step in lipoprotein modification and is, as we report here, evolutionarily and structurally conserved among bacteria and absent from archaea. Our complementation studies show that Lgt is functionally conserved among proteobacteria but differs from Lgt of firmicutes. Lgt from *E. faecalis* and *S. agalactiae* are not functional in *E. coli* and Lgt from *S. aureus* was not detected in *E. coli* lysates even when *lgt* was codon-optimized for expression. Early work, however, showed that plasmid-encoded Lgt of *S. aureus* complements a *lgt*(ts) mutant of *E. coli* (37). The G104S mutation in *lgt*(ts) leads to less efficient Lgt activity, which is reflected by the delay in growth that may be compensated by low amounts of Lgt of *S. aureus* at restrictive temperature.

Subtle differences are observed in growth and viability with Lgt from proteobacteria. Lgt of *S. enterica* serovar Thyphimurium is closely related to *E. coli* Lgt but only fully complements the *E. coli* Lgt depletion strain in absence of Lpp. This observation is somewhat puzzling. *S. enterica* possesses two copies of *lpp* that are both required for virulence (51). Both Lpp-1 and Lpp-2 are highly conserved, as well as to Lpp of *E. coli*, suggesting substrate compatibility between the two bacterial species. Currently, there is no evidence that *S. enterica* possesses multiple copies of *lgt*. Of the Lgt proteins tested from proteobacteria, only Lgt of *N. gonorrhoeae* is unable to compensate loss of viability in the *E. coli* Lgt depletion strain. Currently, the role and composition of the lipid moiety of surface lipoproteins in these processes is unknown.

We showed that the periplasmic head domain is important for Lgt function in cells where high abundant Lpp is present. El Rayes *et al.* recently showed that a disordered linker between the lipid-modified cysteine and the mature folded part of the protein is important for correct sorting to the outer membrane via the Lol machinery in *E. coli* (52). It is therefore likely that the flexible linker domain in preprolipoprotein interacts with the head domain to correctly position itself in the catalytic cavity by its signal peptide.

The arm and head domains are the most diverse, with unique residues located close to arm-2 at the front cleft. For this region, unlike the head domain, a high probability of protein-protein interaction is predicted, which confirms docking studies between Lgt and lipopeptide substrate (31). Shared and essential residues are likely involved in enzyme catalysis, either as catalytic residues or involved in binding of the acyl donor phosphatidylglycerol and preprolipoprotein.

Literature studies, including *in silico* docking experiments with a short lipopeptide substrate, propose an important role for histidine 103 in catalysis (30, 31). Our data show that histidine 103 is poorly conserved and is often substituted by amino acids that cannot serve as a catalytic base in the Lgt reaction. Furthermore, unlike other residues, histidine 103 results in stalling of growth when mutated linked to an aberrant phenotype. Its implication may involve protein substrate binding and correct positioning of preprolipoprotein in the catalytic cavity, while other strictly conserved residues with a more severe phenotype when mutated, are directly involved in catalysis. Our results confirm a conserved role for Y26 in entry of PG substrate, for N146 forming a H-bond with PG upon entry at the front cleft, and for R143 and R239 in positioning the phosphate group of PG in the catalytic site. We propose a Lgt motif where glycine residues provide flexibility to neighbouring catalytic residues that correctly position phosphatidylglycerol and preprolipoprotein in the central cavity. While the X-ray structure and molecular modelling provide insight into the interaction of Lgt and PG (29, 31), a high-resolution structure of Lgt with its preprolipoprotein substrate including a membrane-spanning signal peptide is needed to understand the precise reaction mechanism.

Two inhibitors of *E. coli* Lgt have been reported. Cyclic peptides were synthesised and selected for Lgt binding and tested in an *in vitro* Lgt assay based on cleavage of glycerol-3-phosphate by G3P dehydrogenase after its initial release from a G1P-G3P mixed phosphatidylglycerol (27). Cyclic peptide G2824 containing natural and non-natural amino acids was identified as Lgt inhibitor, also acting on *P. aeruginosa* and *A. baumannii* cells when outer membranes were chemically permeabilised. No effect was observed on *S. aureus* cells, suggesting species specificity. A second report identified a small molecule targeting Lgt using an elegant envelope stress reporter assay in which envelope damage by small molecules leads to release of fluorescent GFP (28). Resistant mutants against MAC-0452936 mapped to Lgt with amino acid substitutions in V109, A37 or G138. V109 is conserved in proteobacteria, except for ε-proteobacteria *H. pylori* and *C. jejuni*, and is located next to unique residue G108 on TM-3 close to essential residues histidine 103, glycine 104 and leucine 106 (29). MAC-0452936 inhibits *E. coli* Lgt activity *in vitro* and overproduction of Lgt reduces its activity. Interestingly, a growth inhibiting effect was observed in *E. coli* in the presence of outer membrane permeabilizer SPR741 (28) but not in *P. aeruginosa* and *A. baumannii*, while increasing antibiotic efficacy was previously observed in *A. baumannii* exposed to SPR741 (53). As for peptide G2428, no inhibitory effect was observed on *S. aureus*.

The functional diversity between Lgt of proteobacteria and firmicutes is likely due to differences in substrate specificity both for the preprolipoprotein and the fatty acid moiety of the phospholipid. A systematic analysis of lipid composition in lipoproteins through lipoproteome identification by mass spectrometry has not been reported for many bacterial species. In general, the lipid moiety reflects the acyl chain composition of membrane phospholipids, although while phospholipids in *H. pylori* contain mainly C14 fatty acids, model lipoproteins are modified by C16:C18 (54). In proteobacteria the main phospholipids are phosphatidylglycerol, cardiolipin (CL) and phosphatidylethanolamine (PE) with C14 to C20 saturated- and unsaturated fatty acids (55). Firmicutes do not possess PE and often have lysyl-PG and galactosyl-(staphylococci) or glucosyl-diacylglycerol (enterococci) as membrane components (56). Branched chain fatty acids are also present in bacilli, including *S. aureus* (57), that influence thickness and fluidity of the membrane and consequently integral membrane protein function. Membranes of *E. faecalis* do not contain branched fatty acids but possess non-saturated carbon chains like *E. coli* (58). This fact might explain the difference we observed in heterologous protein production in *E. coli*, where Lgt of *E. faecalis* is produced but not Lgt from *S. aureus*.

Future biochemical characterisation of Lgt will elucidate how diacylglyceryl transfer takes place and will highlight the basis of substrate specificity between bacteria. Furthermore, the restricted functional conservation between pathogens suggests the potential for developing narrow spectrum rather than broad-spectrum antibiotics targeting Lgt.

## Material and Methods

### Bacterial strains and growth conditions

Unless stated otherwise LB or LB agar (LBA) were used as growth medium. All strains containing the pAM238 plasmid (59) were grown with spectinomycin (50 mg/L). Strain SLEC67 (Δ*lgt*) was grown with 4% L-arabinose to induce wild type *lgt* expression from P*_ara_* (27) and SLEC68 (Δ*lgt*Δ*lpp*) with 0.2% L-arabinose. Strains are listed in Suppl. Table 1. Viability of complementation strains was determined by colony forming units. Selected strains were grown overnight from a single colony in L-arabinose. Cells were washed three times in 1 volume LB without sugar. Finally, they were diluted 1/100 in 5 mL LB in tubes without sugar and grown for 2 hours at 37°C to deplete wild type Lgt. Optical density at 600 nm (OD_600_) was measured and cultures were diluted to an OD_600_ of 0.1. Calibrated cultures were then serial diluted 1/10 in LB to 10^-5^ and 5 μL was spotted onto LBA plates. LBA plates were supplemented with 4%, 0.2% L-arabinose, or 0.2% D-glucose with or without 5 mM IPTG. Plates were incubated overnight at 37°C and imaged on a ChemiDoc imager (Bio-Rad) and colonies counted to determine colony forming unites (CFU/mL).

Complementation of growth in liquid culture was performed by growing elected strains as described above. OD_600_ was recorded and cultures were diluted to an OD_600_ of 0.1 before 100 μL was added to each well of a clear, flat-bottomed 96 well-plate. A volume of 100 μL of LB supplemented with L-arabinose or D-glucose and IPTG was added to a final concentration of 4% or 0.2% L-arabinose, 0.2% D-glucose and 5 mM IPTG. Plates were incubated in a TECAN plate reader at 37°C and regular OD_600_ measurements were taken during 18 hours with shaking between measurements. Results were collected and analysed on GraphPad Prism version 10.

### Plasmid constructions

pCHAP9246 (pAM238-*lgt*^E.c^-*c-myc*_2_) (38) was used as template to generate further IPTG inducible complementation plasmids based on low copy pAM238 vector (Suppl. Table 2). The *c-myc*_2_-tag was replaced with a *flag*_3_-tag (60) as follows. Primers upperFLAG and lowerFLAG (Suppl. Table 3) were annealed in the presence 1 mM MgCl_2_ in 10 mM Tris-HCl buffer pH 7 by slowly cooling down the mixture after heating at 100°C for 5 minutes. The DNA fragment was inserted in XbaI-HindIII digested pCHAP9246. Ligated vector was transformed into chemically competent *E. coli* BW25113 (61) cells and selected for spectinomycin resistance on LBA-Spec50 plates incubated at 37°C overnight. Clones were checked by PCR with M13 F/R primers and sequenced to confirm correct insertion, the plasmid was named SLP14 (pAM238-*lgt*^Ec^*-flag*_3_).

To insert *lgt* genes from different bacterial species SLP14 was used as backbone for gene insertion, through digestion with EcoRI and XbaI. Corresponding *lgt* genes were amplified by PCR from chromosomal DNA with primers listed in Suppl. Table 3 and inserted by Gibson Assembly. To exchange the periplasmic head domain encoding region, plasmids were first linearized by PCR. The primers amplified the vector ‘outwards’ from the flanking regions of the head-domains. The head domain encoding regions were amplified from templates with primers pr140-145 and inserted in the plasmid by Gibson Assembly. All plasmids (Suppl. Table 1) were confirmed by sequencing and the plasmids were transformed into SLEC67 (Δ*lgt*) and SLEC68 (Δ*lgt*Δ*lpp*).

### SDS-PAGE and Western blot detection of Lgt proteins

Cell pellets were resuspended in sample buffer (10% glycerol, 2.5% SDS, 100 mM Tris pH 8, 10 mM DTT, phenol red) to a ratio of 0.01 OD_600_ units/μL. Samples were heated at 100°C for 5 minutes and loaded onto an 12% SDS-PAGE gel. Proteins were transferred onto nitrocellulose membranes and incubated with PonceauS solution, washed with water, and incubated in blocking buffer (5% milk, 0.5% Tween 20 in TBS) for 1 hour at room temperature (RT). The blot was incubated with α-Flag (Sigma) or α-Lgt (custom-made, Eurogentec) antibody solution (0.1% Tween 20 in TBS) for 1 hour at RT. The blot was washed in buffer (0.1% Tween-20 in TBS) and incubated with secondary α-mouse or α-rabbit-HRP antibody (Sigma 1:10,000) in buffer for 1 hour at RT and then washed as described above. The blot was developed using ECL Femto Western blotting chemiluminescence substrate (Pierce) and imaged on a ChemiDoc imager (Bio-Rad) and further analysed using ImageLab software (Bio-Rad).

### Fluorescence microscopy

Cells were harvested corresponding to T_270_ minutes of the growth curve and were fixed in a mixed solutions of formaldehyde (2.8%) and glutaraldehyde (0.04%) for 15 min at room temperature. DNA was stained with Hoechst 2333 (10 μg/mL) and membranes were stained with FM4-64X (0.2 μg/mL). Cells were washed three times in PBS and immobilized on agarose pads (1.2% in PBS) and imaged on a Zeiss Axio Observer fluorescence microscope. Images were analysed using the MicrobeJ version 5.13I plug-in of ImageJ (62, 63), with cell recognition area of 1μm^2^ minimum. Representative images were processed using Adobe Photoshop and Illustrator.

### Sequence identification of Lgt proteins

Species were selected due to their presence as WHO Priority Pathogen (32), ESKAPE pathogen (34), CDC Threat Report pathogen (35), or in a review on global AMR impact (33). From this selection, Lgt protein sequences of representative strains were obtained from the SyntTax database (36). The first strain on the list was selected unless a known reference strain was present. The selected sequence was then compared with nine other strains using the SyntTax web tool to confirm whether it was representative. If variation in synteny or amino acid sequence was observed, up to fifty more strains were analysed. From this collection a suitable representative strain was selected for further analysis. Genes encoding Lgt were searched in the RefSeq genome database using the PFAM (PF01790) definition for the enzyme prolipoprotein phosphatidylglycerol diacylglyceryltransferase as domain using an HMM profile and score threshold suggested by PFAM. Genomes lacking *lgt* or containing multiple *lgt* genes were further analysed by tBlastN and SyntTax (36) to correct sequencing and annotation errors. Next, 179 Lgt protein sequences were retrieved from a selection of 160 bacterial taxa representing all phyla (42) and analysed by multiple sequence alignment. Conservation of the newly defined Lgt motif Y_26_-H_103_-G_104_-G_142_-R_143_-G_145_-N_146_-E_151_-G_154_-Y_235_-R_239_-E_243_-R_246_ (*E. coli* numbering) was extracted from the alignment. The plasmid database WASPS (50) was explored for the presence of extrachromosomal plasmid copies of *lgt*.

### Multiple sequence alignments and phylogenomic analysis of Lgt

Initial sequence alignments were made with Geneious Prime (2024.0.7) using the ClustalO plug-in. Multiple sequence alignment (MSA) of Lgt from 22 strains and 179 Lgt proteins from 160 genomes was carried out using MAFFT v7.481(64) with the option L-INS-I option, and alignment was trimmed using BMGE-1.12 (65) with the default option. Local synteny was determined using SyntTax. Selected strains were queried against Lgt from *E. coli* and a local genome output was generated. Further analysis of predicted operons was conducted via BioCyc (66) and MicrobesOnline (67) operon prediction tools. Finally, Lgt single-gene maximun-likelihood tree was generated with IQ-TREE v2.0.7 (68) and the evolutionary model LG+F+G4 chosen by ModelFinder (69) according to Bayesian Information Criterion (BIC).

### Structural prediction using AlphaFold2

The X-ray crystal structure of Lgt used in this study was PBD 5azc representing the proposed active form of the enzyme in presence of two PG molecules and diacylglycerol (DAG) (29). For structure predictions of Lgt homologues, AlphaFold2 through the ColabFold web tool was used (70). Predicted and solved Lgt structures were compared and essential and conserved residues mapped onto 5azc using ChimeraX (version 1.5). The AF2 model of Lgt from *E. faecalis* was used as representative for Lgt from firmicutes. pIDDT and PAE graphs were obtained to represent confidence and predicted error, respectively.

## Acknowledgements

A Pasteur Paris University Oxford PhD fellowship was awarded to SLG, a PhD fellowship from the Ministère Chargé de l’Enseignement Supérieur et de la Recherche to TR. This project was supported by a Programmes Transversaux de Recherche grant PTR 626-23 from Institut Pasteur Paris, Programme «Bactéries et champignons face aux antibiotiques and antifongiques» (DBF20160635726) and Equipe Fondation pour la Recherche Médicale (FRM) (EQU202403018034). We thank E. Rocha and J. Oberto for initial bioinformatic analysis. Genomic DNA was kindly provided by the Institut Pasteur collection (CRBIP), T. Naas (Ef, Ab), M. Taha (Ng, Hi), S. Dubrac (Sa), A. Firon (Saga), U. Mechtold (Pa), R. Brosch (Mtub) and JM Ghigo (Btheta).

## Supporting information captions

**S1 Fig.1. Synteny of *lgt* containing loci in pathogenic strains.** The Lgt protein sequence of *E. coli* was used in the SyntTax database (36) to search for synteny in the selected species. Identical genes are colour coded. All species have synteny except for *M. tuberculosis* (not shown) but genomic context is not conserved.

**S2 Fig. 2. Confidence metrics and predicted error for Lgt model structures.** AF2 pIDDT and PAE graphs for 11 Lgt-s shown in Fig. 2.

**S3 Fig. 3. Structural position of unique and conserved residues in Lgt of proteobacteria.** (A) Essential residues conserved in 14 Lgt proteins from proteobacteria depicted in Fig. 1. Essential residues are shown in red, non-essential residues in green. Amino acids in black have not been analysed. Unique residues Y30, Y80, M100, S101 and G263 are conserved in proteobacteria and absent from firmicutes and are essential for Lgt function in *E. coli*. (B) Location of essential conserved residues in proteobacteria relative to arm domains (orange) at the front cleft.

**S4 Fig. 4. Sequence alignment and structure prediction of Lgt from *B. thetaiotaomicron*.** (A) Protein sequence alignment of Lgt from *E. coli* and *B. theta* by ClustalO. Among the 16 conserved residues (Fig. 1) 13 are conserved in *B. theta* Lgt (red) and 3 (G142I, Y235I, R246K) are variable (blue). B) AF2 model of Lgt from *B. theta* and (C) corresponding pIDDT and pAE graphs. Arm (orange) and head (pink) domains are shown.

**S5 Fig. 5. Phylogeny of 179 Lgt proteins from 160 genomes representing all phyla highlighting the Lgt motif.** The maximun-likelihood tree was inferred from an alignment of 179 Lgt sequences. The scale bar represents the average number of substitutions per site. Residues of the Lgt signature and active site residues were defined as novel Lgt motif: Y_26_-H_103_-G_104_-G_142_-R_143_-N_145_-N_146_-E_151_-G_154_-Y_235_-R_239_-E_243_-R_246_ by analysis of 179 Lgt proteins from 160 genomes representing all phyla, including *E. coli* K12, *H. pylori* 26695 and *M. tuberculosis* H37Rv as references.

**S6 Table 1. Strains and plasmids used in this study.**

**S7 Table 2. Lgt proteins in ref_seq genomes.**

**S8 Table 3. Primers used in this study**

